# Improving enzyme optimum temperature prediction with resampling strategies and ensemble learning

**DOI:** 10.1101/2020.05.06.081737

**Authors:** Japheth E. Gado, Gregg T. Beckham, Christina M. Payne

## Abstract

Accurate prediction of the optimal catalytic temperature (T_opt_) of enzymes is vital in biotechnology, as enzymes with high T_opt_ values are desired for enhanced reaction rates. Recently, a machine-learning method (TOME) for predicting T_opt_ was developed. TOME was trained on a normally-distributed dataset with a median T_opt_ of 37°C and less than five percent of T_opt_ values above 85°C, limiting the method’s predictive capabilities for thermostable enzymes. Due to the distribution of the training data, the mean squared error on T_opt_ values greater than 85°C is nearly an order of magnitude higher than the error on values between 30 and 50°C. In this study, we apply ensemble learning and resampling strategies that tackle the data imbalance to significantly decrease the error on high T_opt_ values (>85°C) by 60% and increase the overall R^2^ value from 0.527 to 0.632. The revised method, TOMER, and the resampling strategies applied in this work are freely available to other researchers as a Python package on GitHub.

## 1. INTRODUCTION

Enzymes that are stable and active at high temperatures are especially desirable for industrial applications, as they enable biochemical processes to be conducted at higher temperatures yielding faster reaction rates. Hence, researchers have long sought to develop tools for accurate *in silico* prediction of enzyme thermostability. Accordingly, many tools have been developed over the past two decades to predict the enzyme melting temperature (T_m_),^1-3^ the change in thermodynamic stability (ΔΔG) upon point mutations,^4-12^ or the optimal growth temperature (OGT) of the source organism.^13-21^ Unfortunately, for prediction purposes, higher OGT or thermal stability do not necessarily indicate substantial catalytic activity at high temperatures.^22, 23^ Hence, a tool that directly predicts the optimal catalytic temperature (T_opt_) of enzymes is desirable.

Recently, Li *et al.* developed a machine-learning tool, TOME (Temperature Optima for Microorganisms and Enzymes), for predicting the OGT of microorganisms and the T_opt_ of enzymes.^23^ TOME uses a support vector regressor to predict OGT from the dipeptide composition of the proteome, and a random forest regressor to predict T_opt_ from the OGT and the amino acid composition. In predicting OGT, TOME achieved an R^2^ value of 0.88 in cross validation tests, which is superior to other published models.^24, 25^ However, the R^2^ value of T_opt_ prediction was only 0.51, providing impetus for further improvement. More recently,^26^ Li *et al.* incorporated feature engineering to improve the accuracy of T_opt_ prediction. They extracted 5,494 and 5,700 sequence features, using the packages, iFeature and UniRep, respectively.^27, 28^ However, these features did not provide a significant improvement in performance compared to using only the amino acid composition and OGT, even when deep learning was applied. As a result, the authors concluded that more informative features, such as features from the three-dimensional structure, may be necessary to markedly improve T_opt_ prediction performance. Yet, a tool that accurately predicts T_opt_ from sequence-data alone remains valuable to the biotechnology community, since it can be readily applied to the vast number of proteins in the databases that lack structural characterizations.

In this work, we sought to improve the accuracy of T_opt_ prediction, not by customary feature engineering, but by mitigating the adverse impact of the non-uniform distribution of the training data used in the machine learning model. It is recognized that an imbalanced data distribution is highly unfavorable in machine learning problems, as it biases the learning algorithms towards the abundant data regions at the expense of the poorly sampled regions, and, thus, leads to higher error on the rare values and overall sub-optimal model performance.^29-31^ In classification problems, data imbalance has been extensively studied, and numerous techniques for dealing with imbalance problems have been proposed.^32, 33^ These methods are generally classified into three groups: algorithm-level methods, which specifically modify the learning algorithm to address the bias; data-level methods, which resample the data in a preprocessing step to decrease the unevenness of the data; and hybrid methods, which combine both algorithm- and data-level methods.^30, 34^ Data-level methods modify the data distribution primarily by either undersampling the majority class, oversampling the minority class, or a combination of both.^34^ Researchers have developed multiple resampling methods for classification problems such as neighborhood cleaning rule (NCL),^35^ synthetic minority oversampling technique (SMOTE),^36^ selective preprocessing of imbalanced data (SPIDER),^37^ and majority undersampling technique (MUTE).^38^ The combination of resampling strategies with ensemble learning (the integration of the outcomes of multiple base models) has proven remarkably successful in dealing with class imbalance.^34, 39, 40^

On the contrary, less attention has been paid to imbalance in regression problems.^30, 32^ Few methods have been proposed for working with imbalanced distributions in regression domains including: SMOTE for regression (SMOTER),^41^ SMOGN,^42^ meta learning for utility maximization (MetaUtil),^43^ resampled bagging (REBAGG),^44^ and weighted relevance-based combination strategy (WERCS).^45^ In many bioinformatic and cheminformatic supervised-learning regression problems, the data often follows a normal distribution, and the rare extreme values may be more important to the user than the abundant values centered about the median of the distribution. For example, in predicting T_opt_ for practical applications, higher T_opt_ values are generally more relevant since thermostable enzymes are desired for enhanced biochemical reaction rates. Still, a majority of studies do not address the issue of data imbalance,^10, 11, 46, 47^ resulting in models with reduced predictive accuracy at tails of the normal distribution.^32, 45^ Additionally, standard metrics used in assessing regression model performance, such as mean squared error (MSE) and mean absolute deviation (MAD), are heavily biased towards the abundant values centered about the median so that the reported performance fails to capture the poorer performance on rare values at the tails of the distribution.^48^ Consequently, a model could demonstrate excellent performance on non-uniform datasets and, yet, might have little ability to accurately predict extreme values.

In this study, we apply resampling and ensemble methods to enzyme T_opt_ prediction. Our results show that without resampling (i.e., TOME), the error (MSE) in predicting high temperature values (>65°C) was more 500% higher than the error in predicting T_opt_ values centered about the median (30-50°C). By applying resampling strategies alone, without the introduction of new features, we were able to reduce the error on high temperature values (>65°C) by more than 50% and, consequently, increase the overall performance (R^2^) by 20%. We make available the machine-learning tool for improved T_opt_ prediction, TOMER (Temperature Optima for Enzymes with Resampling), through GitHub. We anticipate TOMER will prove valuable in accurately predicting T_opt_ values of industrially-relevant, thermostable enzymes. To facilitate minimizing the impact of data imbalance in other regression applications, we have also provided the resampling strategies employed here as a Python package, resreg (Resampling for Regression).

## 2. METHODS

### 2.1. Dataset and machine learning implementation

The dataset used in training TOMER was obtained from Li *et al.*, consisting of 2,917 enzymes with experimental T_opt_ measurements and OGT data from the BRENDA database.^26, 49^ Throughout this work, all machine learning regressors were trained on the same 21 features used in TOME, which include the frequencies of the 20 amino acids and the OGT. The features were normalized by subtracting the mean and dividing by the standard deviation before fitting the regressors. Machine learning was implemented with the scikit-learn package (v0.21.2)^50^ in Python (v3.6.6).

### 2.2. Evaluation of performance

In evaluating the performance of the regressors, we did not use the conventional *k*-fold cross validation technique. Since the data are normally distributed, randomly splitting the data into folds will result in similarly imbalanced folds and, as a result, the performance metrics (R^2^, MSE) will overly weight the frequent data and will not sufficiently capture the performance at the distribution tails. Hence, we evaluated performance of the regressors on a testing set that was nearly uniformly distributed. A uniform testing set was formed by splitting the entire dataset into five bins based on the target values (T_opt_). Then, 70 samples were randomly selected from each bin to constitute the testing set, with the remaining data forming the training set (**Table 1, Figure 1A**). We selected only 70 samples from each bin so that at least half of the data in the smallest bin (85-120°C) was used in training. This way, 88% of the entire dataset was used in training (2,567 samples) and 12% in testing (350 samples). The dataset was repeatedly split into training and testing sets 50 times, and each time, resampling strategies were applied to the training set before fitting the regressors. The performance on the testing set was measured as an average over the 50 iterations, i.e., Monte Carlo cross validation (MCCV).^51^

**Table 1.** Formation of a uniform testing set by selecting equal samples from five bins.

| Bins | Range (°C) | Samples in bin | Percent of total dataset | Testing size | Training size |
| --- | --- | --- | --- | --- | --- |
| 0-30 | $0 \leq y < 30$ | 461 | 15.8% | 70 | 391 |
| 30-50 | $30 \leq y < 50$ | 1427 | 48.9% | 70 | 1357 |
| 50-65 | $50 \leq y < 65$ | 519 | 17.8% | 70 | 449 |
| 65-85 | $65 \leq y < 85$ | 361 | 12.4% | 70 | 291 |
| 85-120 | $85 \leq y \leq 120$ | 149 | 5.1% | 70 | 79 |
|  | Total | 2,917 | 100% | 350 (12%) | 2,567 (88%) |

**Figure 1.**
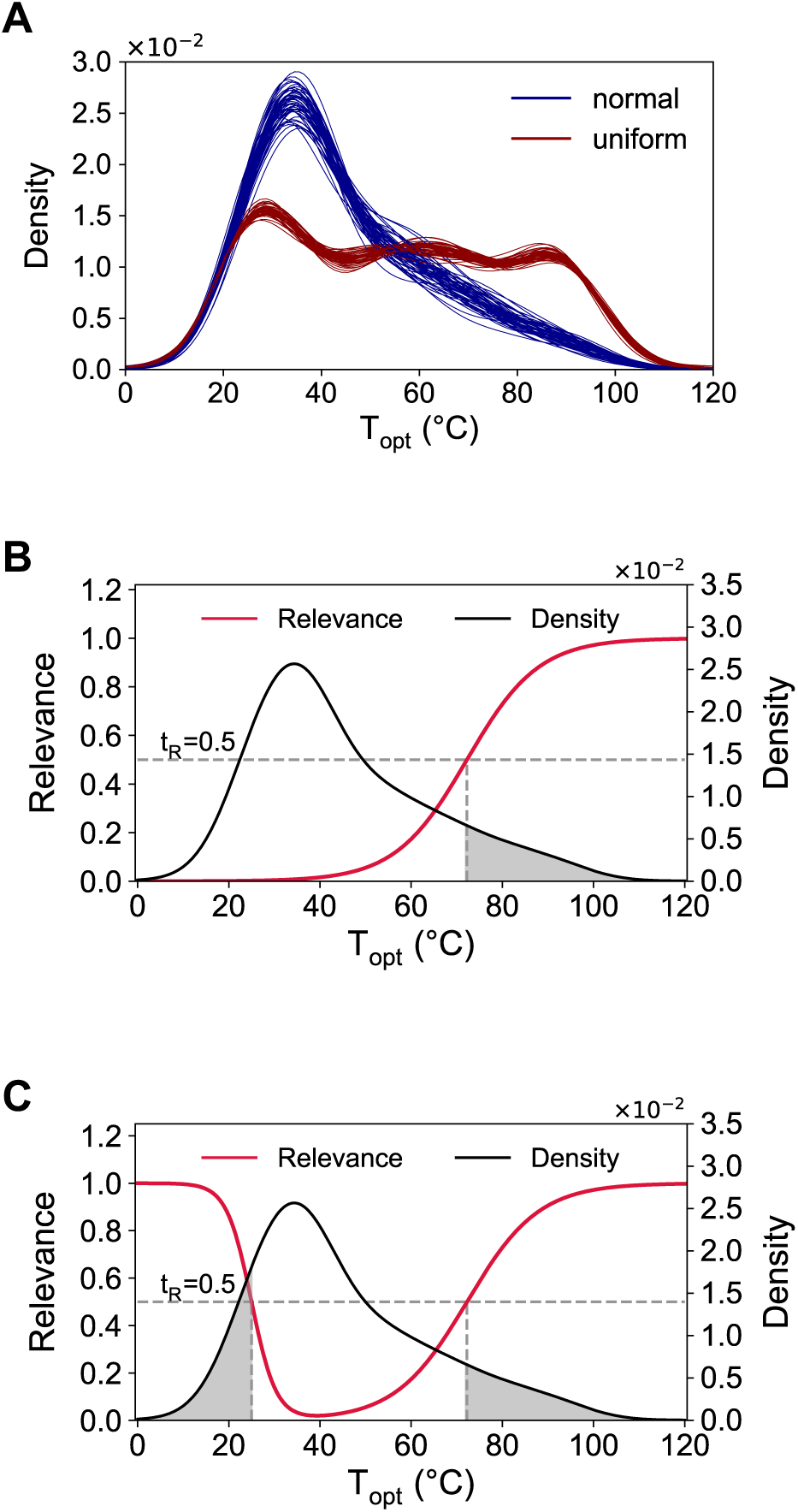
Distribution of T_opt_ values in the dataset of 2,917 proteins. The density plots were derived using a Gaussian kernel density estimation (KDE). (A) Distribution of testing set in 50 iterations of Monte Carlo cross validation. A normally-distributed testing set formed by random selection of 350 samples is shown in blue, and the nearly uniform testing set formed by selecting 70 samples from five bins is shown in red. (B) A one-sided sigmoid relevance function that maps T_opt_ values to relevance values between 0 and 1 (left-hand *y*-axis). By setting the value of *c* in the relevance function (**eq 5**) to the 90^th^ percentile (72.2), T_opt_ values greater than 72.2°C form the rare domain (shaded region) and all other values form the normal domain. The T_opt_ distribution density is shown on the right-hand -axis. (C) A two-sided relevance function mapping T_opt_ values to relevance values between 0 and 1. By setting the values of *c*_*L*_ and *c*_*H*_ in the relevance function (**eq 6**) to be the 10^th^ and 90^th^ percentile (25 and 72.2, respectively), T_opt_ values less than 25°C and greater than 72.2°C form the rare domain, and the complement of the rare domain forms the normal domain. The T_opt_ distribution density is shown on the right-hand *y*-axis.

Four metrics were used to assess the predictive performance. The coefficient of determination (R^2^) on a uniformly-distributed test set was used to assess the overall performance, and was the primary metric for selecting the best resampling strategy. Both real and predicted T_opt_ values were converted to categorical values (0-30 is 1, 30-50 is 2, 50-65 is 3, etc., see **Table 1**), and the Matthew’s correlation coefficient (MCC)^52^ was determined as for a multiclass classification problem.^53^ The mean squared error (MSE) was calculated for each bin to evaluate the variation in the performance across the range of T_opt_ values and to examine the error on rare high values relative to the error on abundant values. Finally, we measured the F_1_ score as a way to assess the predictive performance on high T_opt_ values at the distribution’s tail (≥65°C). The F_1_ score, which is the weighted harmonic mean of precision and recall, is typically a classification performance metric, but has been adapted for regression problems.^41, 48^ For regression, recall and precision has been defined as:^48^

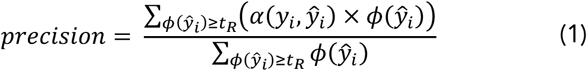

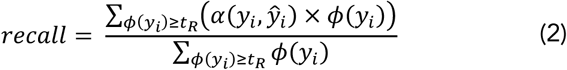

where *y*_*i*_ and *ŷ*_*i*_ are the true and predicted T_opt_ values, respectively; *ϕ*(*ŷ*_*i*_) is the relevance function which maps the target values to a relevance scale from 0 to 1 (discussed below); *t*_*R*_ is the relevance threshold that forms the subdomain of relevant rare values, and *α*(*y*_*i*_, *ŷ*_*i*_) is a function that defines the accuracy of a prediction. Hence, the precision and recall are measures of the predictive accuracy on rare values, weighted by the relevance function. The accuracy function was defined as:^48^

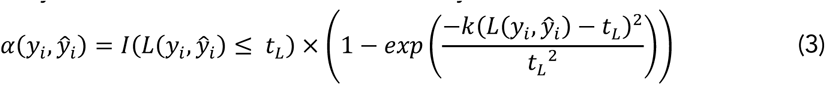

where *L*(*y*_*i*_, *ŷ*_*i*_) is the loss function and is equal to the absolute error of the prediction; *I* is the indicator function, which returns 1 if the absolute error is less than a threshold loss, *t*_*L*_, but zero otherwise; and *k* is an integer that defines the steepness of the accuracy curve. We set *k* to be 10^4^ and *t*_*L*_ to be 5 so that predictions within error limits of 5°C are regarded as accurate. A right-sided relevance function was used, with *t*_*R*_ ≥0.5 for all *y* ≥65 (see **eq 5** and **eq6**), and the F_1_ score was calculated from precision and recall as:

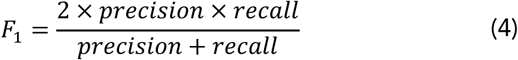

### 2.3. The relevance function

In classification problems, resampling strategies can be readily applied since the target values are clearly divided into discrete classes. Resampling is not as straightforward in regression problems, however, since the target variable is continuous. The concept of a relevance function was introduced in previous works to simplify resampling in regression problems.^41, 48, 54^ The relevance function is a user-defined function that maps the domain of target values to a scale from 0 to 1, where 1 indicates maximum relevance. By specifying a relevance function, *ϕ*(*y*), and a relevance threshold, *t*_*R*_, the domain of target values, *D*, can be split into two sub-domains: a domain of rare values, *D*_*R*_, which is of greater importance to the user, and the domain of normal values, *D*_*N*_ (**Figure 1B and 1C**). Consequently, *D*_*R*_ and *D*_*N*_ can be resampled accordingly.

In this work, we use a sigmoid relevance function defined as:^48^

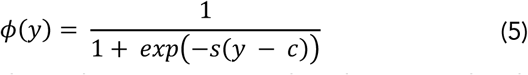

where y is the target variable, and *s* and *c* are constants that determine the shape and center of the sigmoid, respectively. By defining *s* as 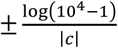, it follows that *s* > 0 implies that *ϕ*(*y*) ≥0.5 for all *y* ≥ *c*, and *s* < 0 implies that *ϕ*(*y*) ≥ 0.5 for all *y* ≤ *c*.^48^ Hence, *c* can be specified so that extreme target values beyond *c* have relevance values above a threshold (*t*_*R*_) of 0.5 and, thus, form the domain of rare values, *D*_*R*_. Otherwise stated, *D*_*R*_ = {*y*: *ϕ*(*y*) ≥ *t*_*R*_} and *D*_*N*_ = {*y*: *ϕ*(*y*) < *t*_*R*_}. Equation 5 is used to determine *ϕ*(*y*) in the case that the rare domain is formed from extreme values at the left or right of the normal distribution (one-sided). For a two-sided rare domain formed from both left and right extremes, we define the relevance function as:

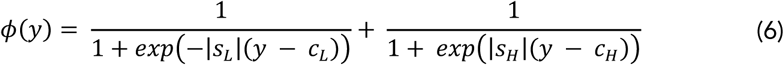

where the subscripts, L and H, indicate low and high extreme values, respectively (**Figure 1B and C**).

### 2.4. Resampling strategies

Having defined a relevance function to split the dataset into a rare and normal domain, we tested several resampling methods that alter the lopsidedness of the rare domain, relative to the normal domain. The resampling methods were applied to the training set to mitigate the adverse effects of the data imbalance, and then a random forest regressor with default settings was fitted to the resampled training set. We adapted and implemented the following resampling strategies in this work: random oversampling (RO), introduction of Gaussian noise (GN), synthetic minority oversampling technique for regression (SMOTER), weighted relevance-based combination strategy (WERCS), and WERCS with Gaussian noise (WERCS-GN).^41, 45^ We give a brief description of these methods below. See the **Supporting Information** for the pseudocode of these methods.

#### 2.4.1. Random oversampling (RO)

With the random oversampling strategy,^45, 55^ the rare values are oversampled by duplicating randomly selected data points, while the normal values are left unchanged. The amount of oversampling is to be specified by the user and can significantly affect the results. Branco *et al.* suggested two automatic methods of oversampling: balance and extreme.^45^ The balance option oversamples the rare domain so that it is equal in size to the normal domain. The extreme option oversamples the rare domain so that the proportion of the size of the rare domain to the size of the normal domain is reversed. For example, if the normal domain is five times larger than the rare domain, the extreme option oversamples the rare domain so that it is five times larger than the normal domain. Here, we introduced a new automatic oversampling method that is intermediate between balance and extreme, which we dub “average”. According to the method selected, the size of the rare domain after oversampling, 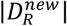, is determined from the size of the rare and normal domain before resampling (|*D*_*R*_| and |*D*_*N*_|, respectively) as follows:

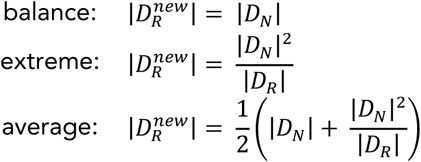

Additionally, the values of *c*_*L*_ and *c*_*H*_, which determine the points at which the target value is split to normal and rare values, can have significant effects on the performance. Hence, we implemented a grid search to determine the optimal combination of hyperparameters for the resampling strategies. We defined the hyperparameter space as *c*_*L*_ ∈ (25, 30, *None*), *c*_*H*_ ∈ (72.2, 60), and *method* ∈ (*balance,average,extreme*) (**Table 2**). The values for *c*_*L*_ correspond to the 10^th^ and 20^th^ percentile of T_opt_, and the values of *c*_*H*_ correspond to the 90^th^ and 80^th^ percentile, respectively. A right-sided rare domain is indicated by *c*_*L*_ = *None* (**Figure 1B and 1C**).

**Table 2.** Hyperparameters of resampling strategies tested with a grid search.

| Strategy | Hyperparameter range |
| --- | --- |
| Random oversampling (RO)<sup>45, 55</sup> | $c_L = (25, 30, \text{None})$, $c_H = (72.2, 60)$, method = (balance, average, extreme) |
| Synthetic minority oversampling technique for regression (SMOTER)<sup>41, 42, 55</sup> | $c_L = (25, 30, \text{None})$, $c_H = (72.2, 60)$, method = (balance, average, extreme), $k = (5, 10, 15)$ |
| Introduction of Gaussian noise (GN)<sup>42, 45, 55</sup> | $c_L = (25, 30, \text{None})$, $c_H = (72.2, 60)$, method = (balance, average, extreme), $\delta = (0.1, 0.5, 1.0)$ |
| Weighted relevance-based combination strategy (WERCS)<sup>45</sup> | $c_L = (25, 30, \text{None})$, $c_H = (72.2, 60)$, over = (0.5, 0.75), under = (0.5, 0.75) |
| Weighted relevance-based combination strategy with introduction of Gaussian noise (WERCS-GN)<sup>42, 45, 55</sup> | $c_L = (25, 30, \text{None})$, $c_H = (72.2, 60)$, over = (0.5, 0.75), under = (0.5, 0.75), $\delta = (0.1, 0.5, 1.0)$ |
| Resampled bagging with random oversampling (BAGG-RO)<sup>44, 55</sup> | $c_L = (25, 30, \text{None})$, $c_H = (72.2, 60)$, method = (balance, variation), $s = (300, 600)$ |
| Resampled bagging with SMOTER (BAGG-SMT)<sup>41, 44, 55</sup> | $c_L = (25, 30, \text{None})$, $c_H = (72.2, 60)$, method = (balance, variation), $k = (5, 10, 15)$, $s = (300, 600)$ |
| Resampled bagging with introduction of Gaussian noise (BAGG-GN)<sup>44, 55</sup> | $c_L = (25, 30, \text{None})$, $c_H = (72.2, 60)$, method = (balance, variation), $\delta = (0.1, 0.5, 1.0)$, $s = (300, 600)$ |
| Resampled bagging with WERCS (BAGG-WR)<sup>44, 45</sup> | $c_L = (25, 30, \text{None})$, $c_H = (72.2, 60)$, over = (0.5, 0.75), under = (0.5, 0.75), $s = (300, 600)$ |
| Resampled bagging with WERCS-GN (BAGG-WRGN)<sup>44, 45, 55</sup> | $c_L = (25, 30, \text{None})$, $c_H = (72.2, 60)$, over = (0.5, 0.75), under = (0.5, 0.75), $\delta = (0.1, 0.5, 1.0)$, $s = (300, 600)$ |

#### 2.4.2. Synthetic minority oversampling technique for regression (SMOTER)

Applying the SMOTER strategy undersamples the normal values and oversamples the rare values by generating synthetic data points through interpolation between each rare value and a random selection of one of its k-nearest neighbors.^41, 42, 55^ The feature vector and target value of a synthetic instance, *X*_2_ and *y*_2_, respectively, are determined as follows:^41^

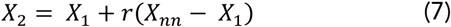

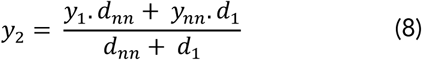

where *X*_1_ is the feature vector of an instance in *D*_*R*_, *X*_*nn*_ is one of *k*-nearest neighbors of *X*_1_, *r* ∈ [0, 1] is a random number, *y*_1_ and *y*_*nn*_ are the target values of *X*_1_ and *X*_*nn*_, respectively, and *d*_1_ and *d*_*nn*_ are the Euclidean distances between *X*_2_ and *X*_1_, and between *X*_2_ and *X*_*nn*_, respectively. The amount of undersampling and oversampling was automatically determined according to the following options:

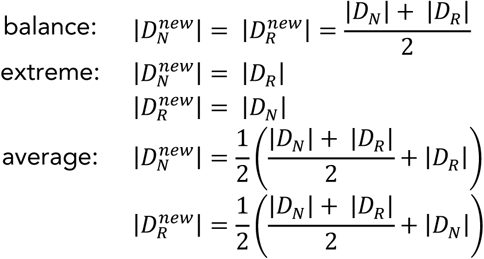

Optimal hyperparameters were similarly determined by a grid search (**Table 2**).

#### 2.4.3. Introduction of Gaussian noise (GN)

The GN strategy is identical to SMOTER in every way except that synthetic points are generated by addition of Gaussian noise rather than interpolation.^42, 45, 55^ Noise based in *N*(0, *δ* × *std*(*a*)) is separately added to each feature and to the target value of a rare instance, where *std*(*a*)) is the standard deviation of the attribute (i.e., feature or target value), and *N* is a user-defined parameter that determines the amplitude of the noise.

#### 2.4.4. Weighted relevance-based combination strategy (WERCS)

Rather than using a relevance threshold to split the data into rare and normal domains as with the previous strategies, the WERCS strategy uses the relevance values as weights to select data points for undersampling and oversampling.^45^ The data are oversampled and then undersampled by selecting instances to be duplicated and instances to be removed, respectively. Selection for oversampling and undersampling is performed using probabilities determined from the relevance function. For each target value in the dataset, *y*_*i*_, we defined the probability that the value is selected for oversampling or undersampling (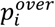 and 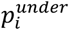, respectively) by **eq 9** and **eq 10**.

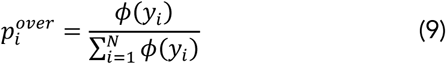

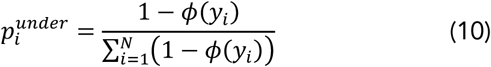

Hence, rare values with higher relevance are more likely to be selected for oversampling and less likely to be selected for undersampling. The amount of oversampling and undersampling are hyperparameters to be specified by the user in percent (*over* and *under*, respectively).

#### 2.4.5. WERCS with Gaussian noise (WERCS-GN)

We modified the WERCS strategy by adding Gaussian noise to the values selected for oversampling by the WERCS strategy. Hence, with WERCS-GN, oversampling is done with synthetic data, instead of by duplicating data points.

### 2.5. Combination of resampling strategies with ensemble learning

Ensemble learning involves training different learners and combining their output to generate a final prediction that is more accurate than the individual learners. Branco *et al.* developed the resampled bagging algorithm (REBAGG) for implementing resampling and bagging in imbalanced regression problems.^44, 56^ In this work, we applied an adaptation of the REBAGG algorithm to the prediction of T_opt_ values, by implementing the resampling methods described previously in the REBAGG algorithm (See the **Supporting Information** for the pseudocode).

First, the dataset is split into rare and normal domains, *D*_*R*_ and *D*_*N*_, using the relevance function, as described previously. Then *N* models are trained on separately resampled bootstrap samples of *s* items from the training dataset. Two modes of the REBAGG method are applied: balance or variation mode. In balance mode, an equal number of samples, ^*s*^/_2_, is randomly drawn from *D*_*R*_ and *D*_*N*_. In the variation mode, however, *p* × *s* samples are drawn from *D*_*R*_, and (1 – *p*) × *s* samples are drawn from *D*_*N*_, where p is a randomly selected number from the set, (1/3, 2/5, 1/2, 3/5, 2/3). Hence, in the variation mode, the *N* models are trained on data that may contain either fewer, equal, or more rare samples than normal samples. If the number of samples to be drawn from *D*_*R*_ is greater than | *D*_*R*_ |, then the extra samples are derived by oversampling the rare domain using RO, SMOTER, or GN, resampling methods as described previously. We refer to the REBAGG method in combination with these resampling methods as BAGG-RO, BAGG-SMT, and BAGG-GN, respectively. A similar combination of REBAGG with WERCS and WERCS-GN (referred to as BAGG-WERCS and BAGG-WRGN) were also implemented. With BAGG-WERCS and BAGG-WRGN, the data are resampled without splitting into rare and normal domains, as in the WERCS and WERCS-GN methods. Then, *s* samples are drawn from the resampled data for training a model in the ensemble. With these resampled bagging strategies, the resampling step is independently repeated for all *m* models with replacement. Finally, each model is applied to the testing set, and the final prediction is determined by averaging the predictions of all *m* models. We used a decision tree regressor with default settings as the base regressor and set *m* to be 100. Other hyperparameters were optimized based on the values shown in **Table 2**.

## 3. RESULTS AND DISCUSSION

### 3.1. Resampling strategies significantly improve predictive performance

In this work, we applied machine learning to predict the T_opt_ of 2,917 enzymes.^23^ The target values follow a normal distribution that creates a problem of data imbalance. Although the T_opt_ values range from 0 to 120°C, about half of the values fall within 30 to 50°C, and high temperature data are scarce (**Table 1**). To deal with this data imbalance, we implemented ten strategies that abate the imbalance by resampling the training data. For each strategy, we tested several hyperparameters with a grid search (**Table 2**) and selected the hyperparameter combination that yielded the highest average R^2^ value on a uniformly-distributed testing set (**Table 3**). Without resampling the training data (i.e., TOME), the average R^2^ value over 50 MCCV iterations was 0.527. However, the best performance of the resampling strategies ranged from 0.567 (RO) to 0.632 (BAGG-RO). Similarly, all resampling strategies yielded significantly higher F_1_ scores (>0.178) and MCC values (>0.235) compared to TOME, which had an F1 score of 0.137 and an MCC score of 0.212 **(Figure 2)**. These results demonstrate that the resampling strategies improve the predictive performance on high T_opt_ values (> 65°C), as illustrated by the higher F1 scores, and lead to superior overall performance, as illustrated by the higher R^2^ and MCC values. It is important to note that some hyperparameter combinations of the resampling strategies led to a reduction in the predictive performance compared to the model that was trained on non-resampled data (TOME) (**Figure S1**). Hence, it is imperative that one test a sufficiently wide range of hyperparameters to determine the optimal hyperparameter combination.

**Table 3:** Best hyperparameter combination for each resampling strategy yielding the highest R^2^ values as determined by a grid search.

| Strategy | Hyperparameter |
| --- | --- |
| RO | $c_L = \text{None}$, $c_H = 60.0$, method=balance |
| SMOTER | $c_L = \text{None}$, $c_H = 60.0$, method=average, $k=10$ |
| GN | $c_L = \text{None}$, $c_H = 72.2$, method=balance, $\delta=0.5$ |
| WERCS | $c_L = \text{None}$, $c_H = 72.2$, over=0.5, under=0.5 |
| WERCS-GN | $c_L = \text{None}$, $c_H = 72.2$, over=0.5, under=0.5, $\delta=0.1$ |
| BAGG-RO | $c_L = \text{None}$, $c_H = 72.2$, method=variation, $s=600$ |
| BAGG-SMT | $c_L = \text{None}$, $c_H = 72.2$, method=variation, $k=5$, $s=600$ |
| BAGG-GN | $c_L = \text{None}$, $c_H = 72.2$, method=variation, $\delta=1.0$, $s=600$ |
| BAGG-WERCS | $c_L = 25.0$, $c_H = 72.2$, over=0.5, under=0.75, $s=600$ |
| BAGG-WRGN | $c_L = 25.0$, $c_H = 72.2$, over=0.75, under=0.75, $\delta=0.1$, $s=600$ |

**Figure 2.**
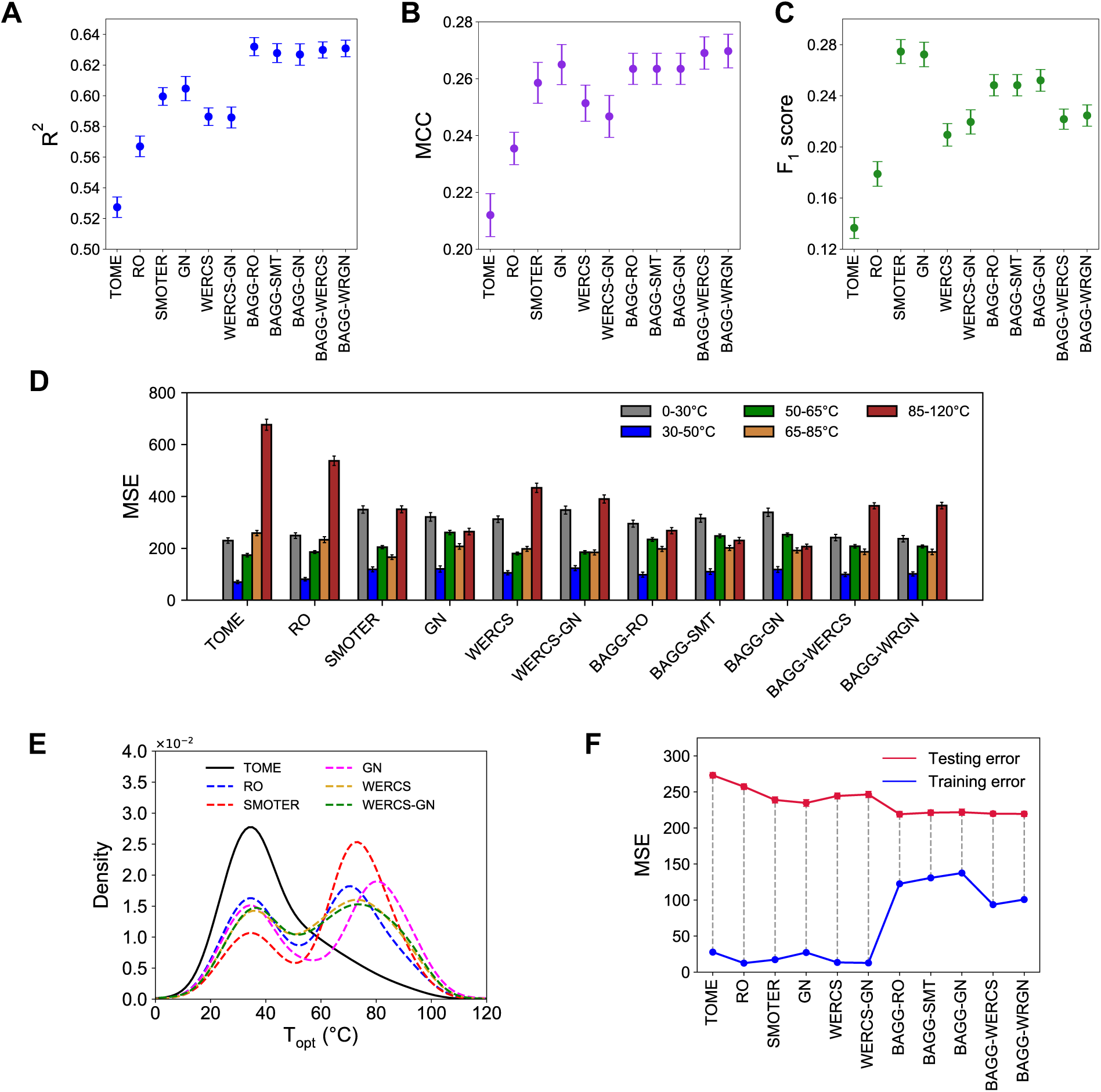
Performance of the resampling strategies. The resampling strategies were applied to the training dataset, regressors were fitted on the resampled data, and the performance was evaluated on a uniformly distributed test set with 50 iterations of Monte Carlo cross validation. Error bars indicate 95% confidence interval of the mean over 50 iterations. (A) Highest R^2^ value of the resampling strategies determined from a grid search of hyperparameter combinations. Combining bagging with the resampling strategies via the REBAGG algorithm outperforms the resampling strategies alone. See **Figure S1** for the performance of all hyperparameter combinations. (B) MCC and (C) F_1_ scores of the best hyperparameter combinations of the resampling strategies, i.e., combinations that yielded the highest R^2^ value. (D) Mean squared error on different ranges of the target values. Without resampling (TOME), the error is highest in the 85-120°C range, but all the resampling strategies significantly reduce this error. The lowest overall error is achieved by the BAGG-RO strategy. (E) Distribution (KDE) of the dataset after applying the resampling methods with optimal hyperparameters. (F) Mean squared error when regressors trained on resampled data are applied to the training set and the testing set. The integration of resampling strategies with bagging decreases the variance as shown by an increase in training error and decrease in testing error.

From the results shown in **Figure 2A-C**, we observed that resampling by simple duplication of rare values, as is done in the random oversampling strategy (RO), led to lower R^2^, F_1_, and MCC values than the strategies that oversample rare values by using the relevance as weights (WERCS, WERCS-GN), or by generating synthetic data through interpolation (SMOTER) or addition of noise (GN, WERCS-GN). However, this trend was not observed when the resampling methods were combined with bagging (BAGG-RO, BAGG-SMT, BAGG-GN, BAGG-WERCS, BAGG-WRGN). We anticipate that duplication performs worse than generating synthetic values because duplication causes the learning algorithms to overfit to the replicated values. Introducing synthetic values, on the other hand, would cause the algorithms to be more general in the rare data region.^32, 36, 57^ Our results indicate that generating synthetic values does not outperform duplication techniques when combined with bagging in the REBAGG strategy, likely because aggregating multiple learners overcomes the overfitting that arises due to replicated values.

Analysis of the MSE as a function of the true T_opt_ values indicates that there is significant variation in the MSE across the range of target values (**Figure 2D**). Without resampling (TOME), the error inversely correlates with the frequency of the data, with lower error in regions of abundant data (30-50°C) and higher error in regions of rare data (0-30°C, 65-120°C). Moreover, error in the 65-85°C and 85-120°C ranges was 3.7 and 9.7 times higher, respectively, than the error in the 30-50°C range. Hence, without resampling, the regressor (TOME) overfits to abundant values and demonstrates inferior performance on high temperature values. In applications that rely on TOME for identifying high T_opt_ enzymes, the large error on high temperature values may lead to misleading results. By applying resampling strategies to the training set, we altered the distribution of the training dataset to prevent the learning algorithm from overfitting to abundant values and to improve performance on rare high temperature values (**Figure 2E**). As **Figure 2D** shows, all the resampling strategies led to a reduction of the error in the high temperature ranges (65-120°C) and an increase of the error in the abundant data range (30-50°C), which indicates a decrease in the overfitting of abundant values. Moreover, the error in the abundant data range is the lowest error for TOME as well as for all the resampling strategies. This suggests that there is an upper limit to the performance gain from resampling rare data, and more experimental data which sample unexplored regions of the rare data space may be necessary for further improvement in performance.

Furthermore, the combination of resampling methods with bagging, such that each base regressor was trained on independently resampled datasets, yielded significantly higher overall performance scores (R^2^ and MCC) than resampling methods alone (**Figure 2A and B**). Other researchers have similarly observed that ensemble learning methods, such as bagging and boosting, considerably enhance the effect of resampling techniques.^34, 39, 58-60^ In this work, the resampling methods without bagging (i.e., RO, SMOTER, GN, WERCS, and WERCS-GN) simply increased the proportion of rare values (**Figure 2E**), which decreased the overfitting of the regressors to abundant data, and, consequently, led to a reduction of both training error and testing error (**Figure 2F**). However, the difference between the testing and training error was substantial, indicating that the regressors were overfitting to the resampled training data (high variance). On the other hand, when the resampling methods were repeatedly applied to multiple decision trees in an ensemble (i.e., the REBAGG strategies) such that each base tree was trained on differently sampled datasets, a much lower testing error and a higher training error was observed. This outcome indicates that the integration of bagging with the resampling methods (i.e., BAGG-RO, BAGG-SMT, BAGG-WERCS, and BAGG-WRGN) reduces the variance of individual regressors and prevents overfitting to the resampled training data, leading to improved generalization.^61^ Moreover, all REBAGG strategies yielded similar overall performance (R^2^ and MCC), which suggests that the specific resampling method applied in the REBAGG strategy had little effect on the overall performance. The BAGG-RO strategy led to the highest R^2^ value of 0.632 and the lowest MSE of 218.6.

### 3.2. Effect of base learners on ensemble performance

We examined the influence of different base learners in the BAGG-RO ensemble to assess whether further performance enhancement could be attained. Using the optimal resampling hyperparameters determined with decision trees (**Table 3**), we applied four additional base regressors in the BAGG-RO ensemble: support vector regressor (SVR), *k*-neighbor regressor (KNR), elastic net (ENET) regressor, and Bayesian ridge regressor (BAYR). For each of these regressors, we used a grid search to determine optimal hyperparameters that yielded the best R^2^ value **(Table 4)**, and the performance was measured as an average over 50 MCCV iterations. The results indicate that, although each alternative regressor outperformed TOME, the decision tree base regressor yielded the highest R^2^ value and lowest overall MSE. Interestingly, the decision tree regressor showed the lowest F_1_ score (**Figure 3**). These results suggest that, while other regressors possibly perform better on high temperature values, tree-based regressors exhibit the best overall performance in predicting T_opt_ values from amino acid composition and OGT.^23, 26^

**Table 4:** Hyperparameters for base learners in BAGG-RO ensemble.

| Base learner | Hyperparameter range | Optimal hyperparameters |
| --- | --- | --- |
| Support vector regressor (SVR) | C = [10<sup>-3</sup>, 10<sup>-2</sup>, 10<sup>-1</sup>, 10<sup>0</sup>, 10<sup>1</sup>, 10<sup>2</sup>],<br>gamma=[10<sup>-3</sup>, 10<sup>-2</sup>, 10<sup>-1</sup>, 10<sup>0</sup>, 10<sup>1</sup>, 10<sup>2</sup>] | C=10<sup>2</sup>,<br>gamma=10<sup>-2</sup> |
| k-neighbor regressor (KNR) | k=[3, 5, 7, 10, 15, 20, 30] | k=3 |
| Elastic net regressor (ENET) | alpha=[10<sup>-3</sup>, 10<sup>-2</sup>, 10<sup>-1</sup>, 10<sup>0</sup>, 10<sup>1</sup>, 10<sup>2</sup>] | alpha=10<sup>-2</sup> |
| Bayesian ridge regressor (BAYR) | None (default) | None (default) |

**Figure 3.**
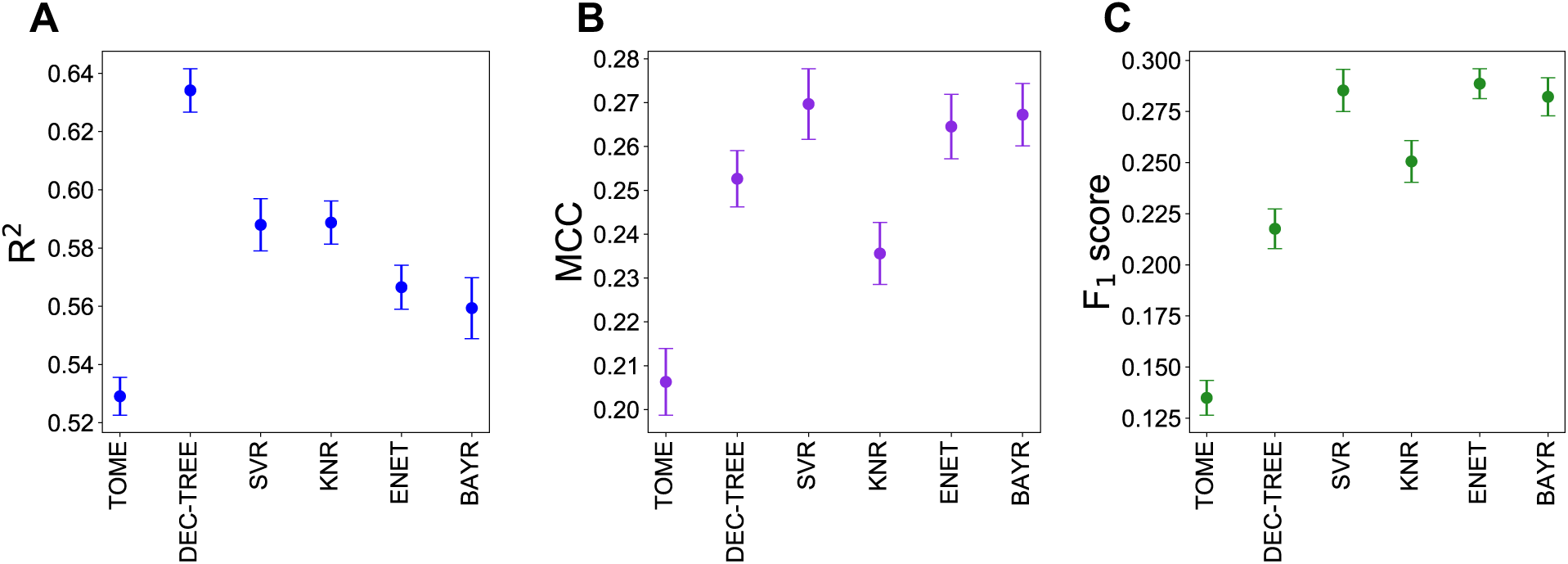
Performance of BAGG-RO ensemble with different base learners. The optimal hyperparameters for the base learners were determined by a grid search. (A) Highest R^2^ value achieved for different base learners in the BAGG-RO ensemble. (B) Matthew’s correlation coefficient and (C) F_1_ scores of BAGG-RO strategy with different base learners using the optimal hyperparameters, i.e., hyperparameters that yielded the highest R^2^ value.

### 3.3. Final model, data and code availability

We identified the BAGG-RO strategy with decision tree base learners as the optimal resampling strategy for predicting enzyme optimum temperatures across the entire range of experimental T_opt_ values because it led to the highest R^2^ value and lowest overall MSE. A final model was prepared by applying the BAGG-RO resampling strategy with optimal hyperparameters **(Table 3)** to the entire dataset of 2,917 proteins. The final model is available to researchers as a Python package, TOMER (Temperature Optima for Enzymes with Resampling), on the Python package index, http://pypi.org/project/tomer/ with the source code publicly available at http://github.com/jafetgado/tomer/. Compared to TOME, TOMER provides a 20% improvement in the overall predictive performance (R^2^), and a 25% and 60% decrease in MSE on T_opt_ values in the 65-85°C and 85-120°C ranges, respectively. All data and code used and produced in this study are available at https://github.com/jafetgado/tomerdesign/. We have also prepared a Python package, resreg (resampling for regression), for applying the resampling strategies discussed in this work to other regression problems. It is available on the Python repository, http://pypi.org/project/resreg, with the source code at http://github.com/jafetgado/resreg.

## 4. CONCLUSIONS

In this study, we applied resampling strategies to improve the performance of predicting enzyme optimum temperatures with machine learning. The resampling strategies were implemented to modify the imbalanced distribution of the training set and improve performance on regions with sparse data. Compared with TOME, which at the time of this study is the only available machine-learning tool for predicting enzyme optimum temperatures, our method (TOMER) yields a significant improvement in predictive accuracy, particularly in the thermophilic regimes. We expect that TOMER will find useful application in high-throughput prospecting of enzymes that are both stable and active at high temperatures. TOMER requires the user to provide the amino acid sequence of the enzyme and the OGT of the source organism. If the OGT is unknown, it may be predicted using TOME.^23^ For future considerations, the incorporation of higher-level features or the addition of more experimental data may prove useful strategies for further improving the performance of TOMER. Ultimately, this study highlights the critical need to consider data imbalance in regression problems, especially when the rare, extreme data range is of greater scientific interest than the abundant data region. We anticipate that our Python tool for readily implementing resampling strategies in regression problems (resreg) will be a valuable resource for other researchers in dealing with the challenges of data imbalance.

## ASSOCIATED CONTENT

### Supporting Information

Figure S1, performance of resampling strategies for all hyperparameter combinations Algorithms (pseudocode) for the RO, SMOTER, GN, WERCS, WERCS-GN, and REBAGG resampling strategies.

## AUTHOR INFORMATION

### Notes

The authors declare no competing financial interest.

## ACKNOWLEDGMENTS

This work was partially supported by the National Science Foundation (CBET-1552355 to CMP in support of JEG). Support for GTB was provided by the U.S. Department of Energy, Office of Energy Efficiency and Renewable Energy, Advanced Manufacturing Office (AMO) and Bioenergy Technologies Office (BETO). This work was performed as part of the BOTTLE™ Consortium and was supported by AMO and BETO under contract no. DE-AC36-08GO28308 with the National Renewable Energy Laboratory, operated by Alliance for Sustainable Energy, LLC. This material is also based upon work supported by (while CMP is serving at) the NSF. Any opinion, findings, and conclusions or recommendations expressed in this material are those of the authors and do not necessarily reflect the views of the NSF. The authors also thank Dr. Peter St. John (NREL) and Dr. Brent Harrison (University of Kentucky) for a critical reading of the manuscript.

---

**Figure:**
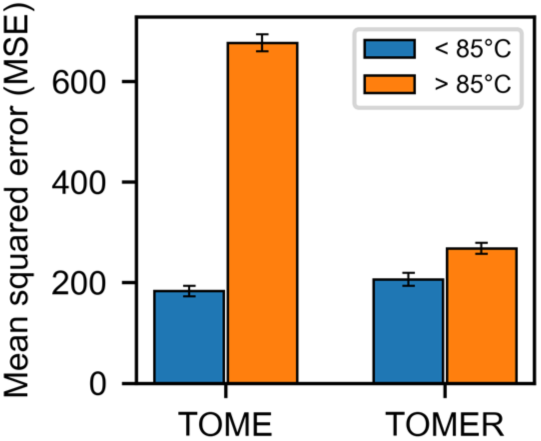
For Table of Contents Only

