## Supporting Information for "Improving enzyme optimum temperature prediction with resampling strategies and ensemble learning"

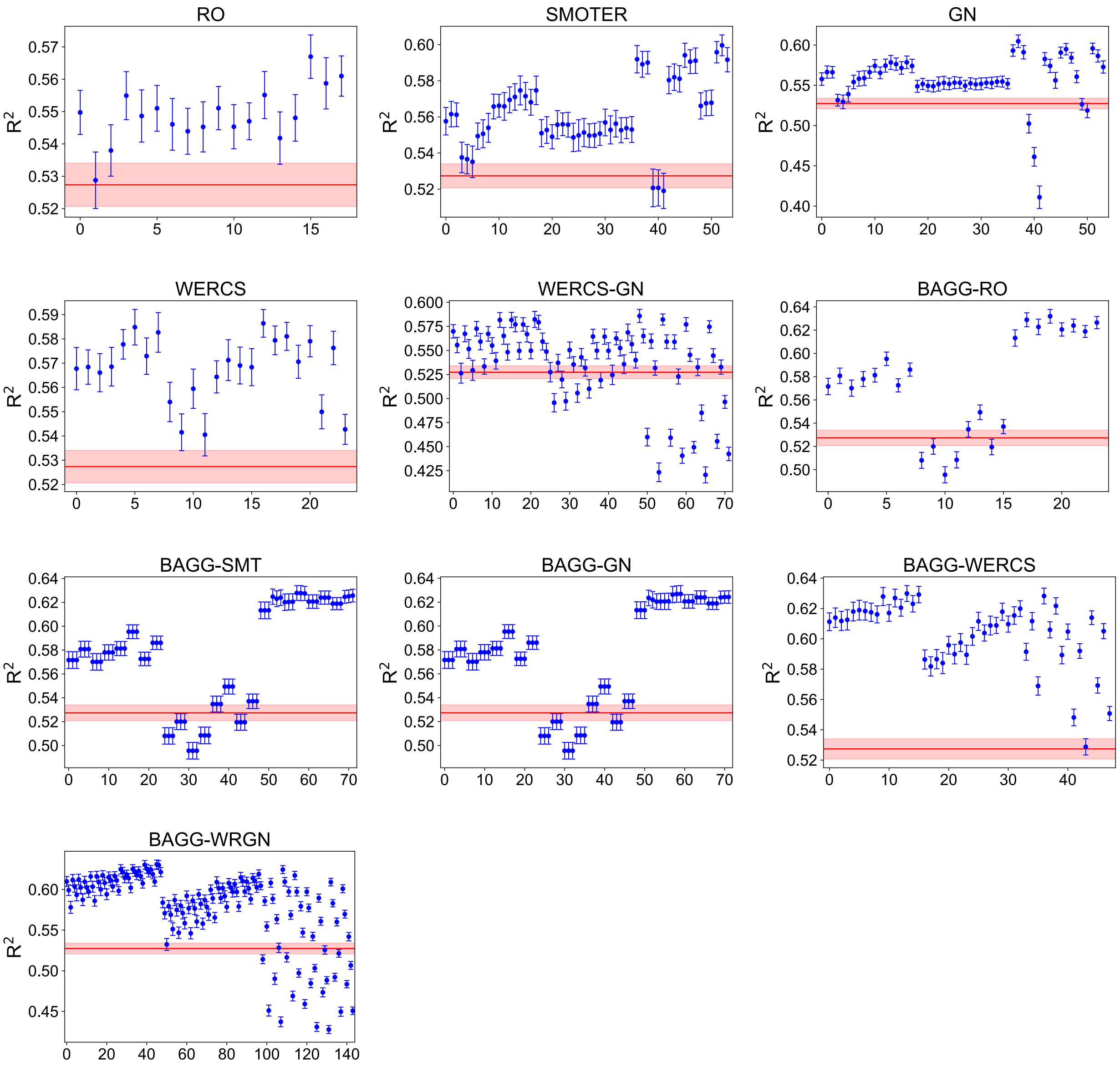


**Figure S1:** Performance ($R^2$) of resampling strategies for all hyperparameter combinations. The *x*-axes indicate different hyperparameter combinations in a grid search. Error bars indicate 95% confidence interval of the mean determined from averaging results over 50 Monte Carlo cross validation repetitions. The red line represents the baseline performance obtained when a random forest regressor is applied to the dataset without resampling (TOME) and the shaded region around the red line indicate the 95% confidence interval. From the figure, it is apparent that some hyperparameter combinations lead to inferior performance relative to TOME. The BAGG-RO strategy with hyperparameters: $C_L$=None, $C_H$=72.2, *s*=600, and *method*=variation, yielded the highest $R^2$ value of 0.632.

### Pseudocode for resampling strategies as applied in this work

**Algorithm 1**: Random oversampling algorithm (RO)[1, 2]

**Input**:

*D (X, y)* – dataset with features, *X*, and continuous target value, *y*
*R* – relevance values for corresponding target values
$t_R$ – relevance threshold
*over* – oversampling percent

**Output**: *newD* – resampled dataset

*rareD* ← instances in *D* with relevance ≥ $t_R$
*normD* ← instances in *D* with relevance < $t_R$
*addSize* ← *over* × |*rareD*| //`or determine addSize by balance, extreme, or average methods`
newRareD ← rareD ∪ randomly selected *addSize* instances from *rareD*
*newD* ← *normD* ∪ *newRareD*

**return** *newD*

**Algorithm 2**: SMOTER algorithm[2-4]

**Input**:

*D* (X, y) – dataset with features, *X*, and continuous target value, *y*
*R* – relevance values for corresponding target values
$t_R$ – relevance threshold
*over* – oversampling percent
*under* – undersampling percent
*k* – number of nearest neighbors

**Output**: *newD* – resampled dataset

*normD* ← instances in *D* with relevance < $t_R$
*rareD* ← instances in *D* with relevance ≥ $t_R$
*lessSize* ← *under* × |*normD*|
*addSize* ← *over* × |*rareD*| //`or determine addSize and lessSize by balance, extreme, or average methods`
*rareDL* ← instances in *rareD* with *y* < median(*y*)
*rareDH* ← instances in *rareD* with *y* ≥ median(*y*)
*addSizeL* ← *addSize* × $\frac{|rareDL|}{|rareD|}$
*addSizeH* ← *addSize* × $\frac{|rareDH|}{|rareD|}$
*nns* ← get k-nearest neighbors of all instances in *rareD*
*newRareDL* ← generate *addSizeL* instances by interpolating between points in *rareDL* and a random selection of one of the k-nearest neighbors
*newRareDH* ← generate *addSizeH* instances by interpolating between points in *rareDH* and a random selection of one of k-nearest neighbors
*newRareD* ← *newRareDL* ∪ *newRareDH* ∪ *rareD*
*newNormD* ← *normD* \ random selection of *lessSize* instances from *normD* //`undersampling`
*newD* ← *newNormD* ∪ *newRareD*

**return** *newD*

/ /

/ 

**Algorithm 3**: Introduction of Gaussian noise algorithm (GN)[1, 2, 4]

**Input**:

*D* (X, y) – dataset with features, *X,* and continuous target value, *y*
*R* – relevance values for corresponding target values
$t_R$ – relevance threshold
*over* – oversampling percent
*under* – undersampling percent
$\delta$ – magnitude of Gaussian noise

**Output**: *newD* – resampled dataset

```
normD ← instances in D with relevance < t_R
rareD ← instances in D with relevance ≥ t_R
lessSize ← under × |normD|
addSize ← over × |rareD|   //or determine addSize and lessSize by balance, extreme, or average methods
newRareD ← random selection of addsize instances from rareD

foreach case in newRareD do //add Gaussian noise to each case
        foreach a in X ∪ y do
                case[a] = case[a] + N(0, δ×std(a))
        end
end
newNormD ← normD \ random selection of lessSize instances from normD    //undersampling
newD ← newNormD ∪ newRareD

return newD
```

**Algorithm 4**: Weighted relevance combination strategy (WERCS and WERCS-GN) algorithm[1]

**Input**:

*D* (X, y) – dataset with features, *X,* and continuous target value, *y*
*R* – relevance values for corresponding target values
*over* – oversampling percent
*under* – undersampling percent
$\delta$ – magnitude of Gaussian noise

**Output**: *newD* – resampled dataset

*underSize* ← *under* × |*D*|
*overSize* ← *over* × |*D*|
*pOver* ← $\left\{\frac{r_i}{\sum r_i} \mid r_i \in relevance\right\}$
*pUnder* ← $\left\{\frac{1-r_i}{\sum 1-r_i} \mid r_i \in relevance\right\}$

```
underD ← sample underSize instances from D with pUnder weights
newD ←D \ underD    //undersample
overD ← sample overSize instances from D with pOver weights

if method is WERCS-GN do   //add Gaussian noise
        foreach case in overD do
                foreach a in X ∪ y do
                        case[a] = case[a] + N(0, δ×std(a))
                end
        end
end
newD ←newD ∪ overD

return newD
```

/

/

**Algorithm 5**: Resampled bagging (REBAGG) algorithm[1, 2, 4, 5]

```
Input:
        D (X, y) – dataset with features, X, and continuous target value, y
        m – number of models in ensemble
        s – size of bootstrap samples
        reg – regressor with specified parameters
        R – relevance values for corresponding target values
        tR – relevance threshold
Output: yPred – predictions for a test set (XTest)
//Fitting the ensemble
for i ←1 to m do
        If sample method is RO, SMOTER, or GN do
            normD ← instances in D with relevance < tR
            rareD ← instances in D with relevance ≥ tR
            if size method is balance do
                rareSize = normSize = s/2
            elseif size method is variation do
                rareSize = s × r ∈ [1/3, 2/5, 1/2, 3/5, 2/3]    \\ r is a random number
                normSize = s – rareSize
            end
            If sample method is RO do
                newRareD ← randomly sample rareSize instances from rareD
            elseif sample method is SMOTER do
                newRareD ← sample rareSize instances from rareD with SMOTER algorithm  // by interpolation
            elseif sample method is GN do
                newRareD ← sample rareSize instances from rareD with GN algorithm  // add Gaussian noise
            end
            newNormD ← randomly sample normSize instances from normD
            newD ← newRareD ∪ newNormD
        elseif sample method is WERCS or WERCS-GN do
            newD ← oversample and undersample D with relevance as weights
            if sample method is WERCS-GN do
                newD ← add Gaussian noise to newD with GN algorithm
            end
        end
        baseModel_i ← model trained with regressor reg on resampled data newD
end
// Applying the ensemble to obtain predictions
for X_k in XTest do
        yPred_k ← (Σ_{i=1}^{m} prediction of baseModel_i on X_k) / m
end
return yPred
```
